# Artificial Selection on *Storage Protein 1* Contributes to Increase of Hatchability during Silkworm Domestication

**DOI:** 10.1101/388843

**Authors:** Yanan Zhu, Lizhi Wang, Cencen Li, Yong Cui, Man Wang, Yongjian Lin, Ruoping Zhao, Wen Wang, Hui Xiang

## Abstract

Like other domesticates, efficient utilization of nitrogen resource is also important for the domestic insect, the silkworm. Deciphering how artificial selection act on silkworm genome for improved utilization of nitrogen resource and further human-favored domestication traits will provide unique cues from the insect scenario for understanding general rules of Darwin’s evolutionary theory on domestication. Storage proteins (SP), which belong to a hemocyanin superfamily, basically serve as a source of amino acids and nitrogen during metamorphosis and reproduction in insects. Here through genomic search and further screening of artificial selection signature on silkworm SPs, we discovered a candidate domestication gene, i.e. the methionine-rich storage protein1 (*SP1*), which is uniquely diverged from the others and showed increased expression in the ova of domestic silkworms. Knockout of *SP1* via CRISPR/Cas9 approach resulted in dramatic decrease in egg hatchability, without obvious impact on egg production, which was similar to the case in the wild silkworm compared with domestic one. Larval development or metamorphosis were not affected by *SP1* knockout. Comprehensive ova comparative transcriptomes indicated a general repression of gene expression, specifically vitellogenin, chorion proteins and structural component proteins in the extracellular matrix (ECM)-interaction pathway, as well as enzymes in folate biosynthesis, in both the mutant and the wild silkworm with the mutated allele, compared to the wild type domestic silkworm. Wild silkworms with the wild allele also showed generally down-regulated expression of genes enriched in structural constituent of ribosome and amide and peptide biosynthesis. This study exemplified a novel case that artificial selection could directly act on nitrogen resource protein to affect egg nutrient and eggshell formation, and activate ribosome for improved biosynthesis and increased hatchability during domestication. The findings shed new light on both understanding of artificial selection and silkworm breeding from the angle of nitrogen and amino acid resource.

**Author summary:** Like other domesticates, nitrogen resource is also important for the domestic insect, the silkworm. Deciphering how artificial selection act on silkworm genome for improved utilization of nitrogen resource and further human-favored domestication traits, will provide unique cues from insect scenario, for understanding general rules of Darwin’s evolutionary theory. However, mechanism of domestication in the silkworm is largely unknown to date. Here we focused on one important nitrogen resource, i.e, the storage proteins (SP). We discovered that the methionine-rich storage protein1 (*SP1*) which is divergent from the other SPs are the only target of the artificial selection. We proposed based on functional evidence together with the key findings of comprehensive comparative transcriptome, that artificial selection, on one hand favored higher expression of *SP1* in the domestic silkworm, which would subsequently up-regulate the genes or pathways vital for egg development and eggshell formation. On the other hand, artificial selection consistently favored activated ribosome activities and improved amide and peptide biosynthesis and in the ova, as it might act in the silk gland for increased silk-cocoon yield. We here exemplified a novel case that artificial selection could directly act on nitrogen resource protein for human desired domestication trait.

## Introduction

The silkworm Bombyx *mori* is the only fully domestic insect species, which originated from its wild ancestor *B. mandarina* about 5000 years ago. During this process, the domestic silkworm evolved rapidly under human-preferred selection. Deciphering how artificial selection act on silkworm genome for human-favored domestication traits, will provide unique cues from insect scenario, for understanding general rules of Darwin’s evolutionary theory. Recently, by population and evolutionary genomic analyses on domestic and wild silkworm individuals, we recently found that nitrogen and amino acid metabolism pathways and specifically genes in glutamate and aspartate metabolism, were under artificial selection and could affect the metamorphosis and cocoon yield [1]. These findings suggest that, like domestic plants and animals, domestic silkworms also tend to have efficient utilization of nitrogen resources to adapt to human-preference [1–3]. Besides the glutamate and aspartate metabolism which is an ammonia re-assimilated system[4], we further wonder whether other kind of nitrogen resources are also affected by artificial selection. If this is the case, how they contributed to silkworm phenotypic changes during domestication.

Insect storage proteins (SP) are another important source of amino acids and nitrogen, which belong to a special conserved arthropod hemocyanin superfamily [5]. Most insects have at least two main kinds of storage proteins, i.e, arylphorin and methionine-rich storage proteins and some species have other non-typical SPs [6]. *SPs* have been cloned or predicted in many insect species, including Lepidoptera moths and butterflies [7–10]. Insect SPs are suspected to serves as a source of amino acids and nitrogen the for pupae and adults during metamorphosis and reproduction [11], however solid functional evidence on its biological significance is rather few [9, 12]. In plant, storage proteins are mainly reserved in seeds, together with other nutrient such as oil and starch, to supply energy for seed germination, growth [13, 14]. Especially in crops, seed SPs function in providing energy for humans and animals and they are of great interest and target for breeding and improvement [13–15].

In the domestic insect, the silkworm, previous studies preliminarily characterized three SPs mainly based on cues of gene or protein expression pattern [7, 16–18]. Especially the methionine-rich SP1 was implied to contribute to adult female characters [7] and related to synthesis of vitellogenin (Vg), the precursor of yolk protein [16]. SP2 coupled with SP3 form a heterohexamer and has the inhibitory effects on cell apoptosis [17, 18]. Whether SPs are also important in the silkworm domestication, as observed in domesticated plants, given the importance of nitrogen supply in silkworm domestication, pends deep exploration of their biological and evolutionary significances.

Development of genomics and genome-editing techniques provide tools for efficiently decipher the evolutionary and functional significances of interested genes [19, 20]. Here in this study, we conducted a genome-wild identification of silkworm SPs and taking advantages of the genomic data resource of a batch of representative domestic and wild silkworms [1], we conducted selection signature screening of these silkworm SPs followed by functional verification via CRISPR/Cas9 knockout system and comprehensive comparative ova transcriptomes of the wild type and mutant silkworms as well as domestic and wild silkworms. The findings here suggested that artificial selection on *SP1* contribute to increased production during silkworm domestication, possibly by upregulation of vitellogenin and egg development and eggshell formation, to promote egg hatchability during domestication. These findings provide a novel case with functional evidence and figure out a frame of regulation on a silkworm domestication gene, which illuminated that artificial on nitrogen and amino acid supply will be also required for improved silkworm reproduction.

## Results

### SP1 *is diverged from the other* SPs *and is the only one targeted by artificial selection*

Totally, we identified 7 *SPs* in the silkworm genome by blast search. SP1 showed the dramatically highest methionine content (10.98%) (Table 1). Phylogenetic analysis showed that *SP1* was one distinct clade whereas the others were in another one, indicating an obvious divergence between *SP1* and the others (Fig 1A). *SP1* is located in Chromosome 23 and the others are clustered in Chromosome 3, suggesting possible tandem duplication events during evolution. Interestingly, by screening of artificial selection signatures on the genomic region bearing *SP1* and the other *SPs* respectively, we detected strong signatures in the *SP1* region (see material and methods) (Fig 1B and 1C). Furthermore, we detected strong differentiation in allelic frequency in the upstream of *SP1* (Fig 1D). Correspondingly, *SP1* were differentially expressed in the ova between domestic and wild silkworms, with higher expression in the domestic one (Fig 1E). We also detected 11 SNPs that caused amino acid changes in the coding sequence of the gene (S1 Fig), despite that the biological significance of these SNPs needs further evaluation. These results suggest that artificial selection on *SP1* during silkworm domestication might affect the function of this gene in domestic silkworms, at least in terms of gene expression.

**Table 1.** Basic information and methionine composition of silkworm SPs.

|  | **Protein length**<br>**(KD)** | **Protein weight**<br>**(No. amino acid)** | **No. Met** | **Proportion of**<br>**Met (%)** |
| --- | --- | --- | --- | --- |
| **SP1** | 87.25 | 747 | 82 | **10.98** |
| **SSP2** | 83.45 | 703 | 23 | 3.27 |
| **SP2** | 83.46 | 704 | 27 | 3.84 |
| **SP3** | 82.85 | 696 | 22 | 3.16 |
| **Bmor_9524** | 87.39 | 743 | 49 | 6.59 |
| **Bmor_9525** | 99.02 | 866 | 18 | 2.08 |
| **Bmor_9527** | 77.61 | 651 | 20 | 3.07 |
| **Bmor_9880** | 47.42 | 441 | 3 | 0.73 |

**Figure 1.**
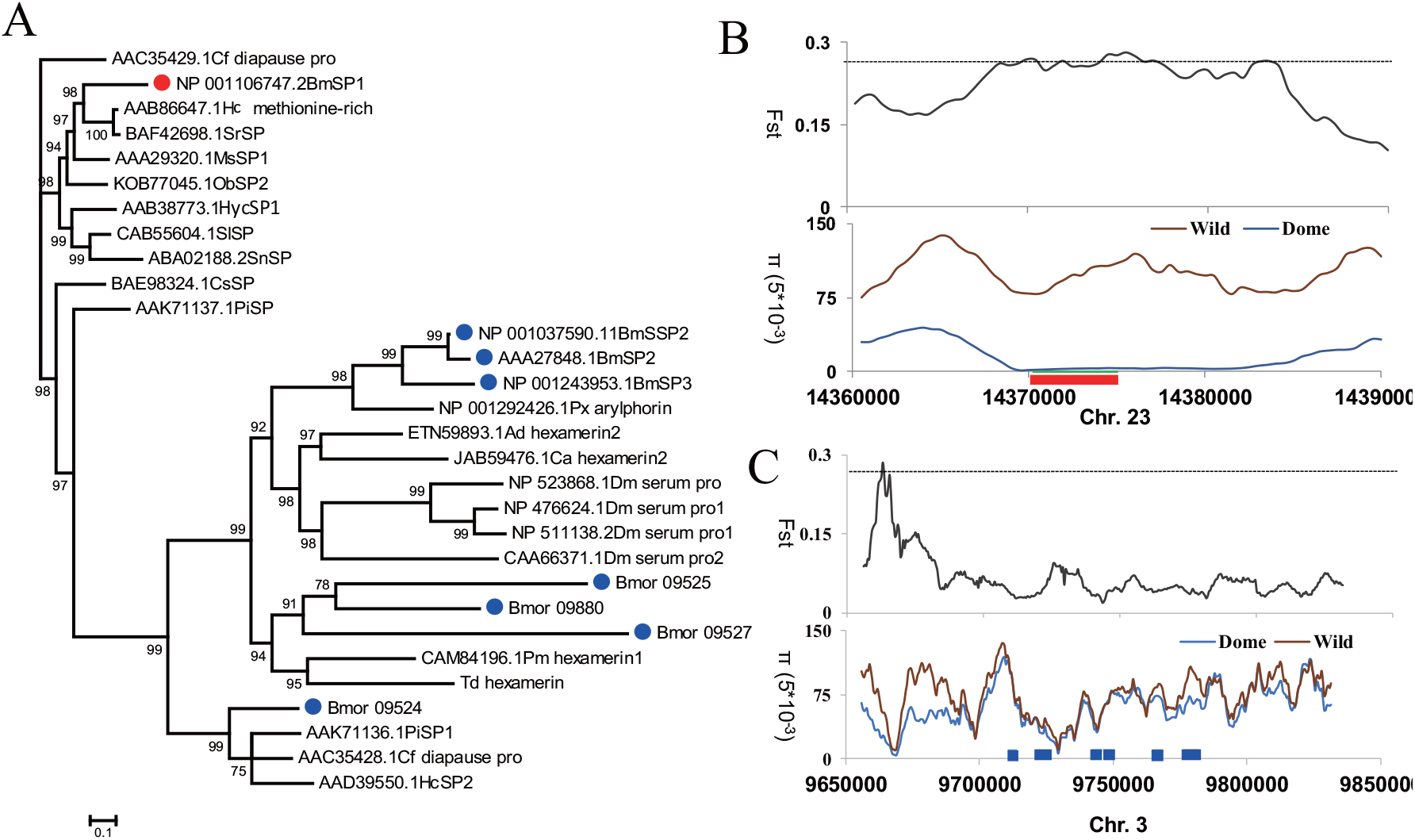
Molecular evolution of silkworm SPs. **(A)** Phylogenetic tree of insect Storage proteins based on Bayesian inference analyses. Full-length amino acid sequences were aligned to generate a phylogenetic tree. Bayesian posterior probability was shown for each node. Cf, *Choristoneura fumiferana*; Bm, *Bombyx mori*; Hc, *Hyalophora cecropia*; Sr, *Samia ricini*; Ms, *Manduca sexta*; Ob, *Operophtera brumata*; Hyc, *Hyphantria cunea*; Sl, *Spodoptera litura*; Sn, *Sesamia nonagrioides;* Cs, *Chilo suppressalis*; Pi, *Plodia interpunctella*; Px, *Plutella xylostella*; Ad, *Anopheles darlingi;* Ca, *Corethrella appendiculata;* Dm, *Drosophila melanogaster;* Pm, *Perla marginata*; Td, *Thermobia domestica*. Accession number for each protein is indicated ahead of abbrevation of each species. **(B-C)**. Selection signatures of the silkworm SPs. Signature index— Population divergence coefficient (Fst) between the Chinese local trimoulting(CHN_L_M3) domestic silkworm group and the wild silkworms and nucleic acid diversity (π) in the silkworm populaton is shown along the genomic regions covering the *SP* genes. Dashed lines represent the top 1% Fst cutoff. The red square represents *SP1* region which is located in Chromosome 23 and the blue squares represents the other *SPs* which are clustered in Chromosome 3. **(D)** Plotting of frequency of reference genotype for each SNP position in the upstream 2 kb region of *SP1*, indicating many mutant alleles with fairly high allelic frequency in the wild silkworm population. **(E)** Expression level indicated as FPKM of *SP1* in the ova of domestic and wild silkworms. The three data pot indicate the highest value with 95% confidence, the average and the lowest value with 95% confidence, respectively. ***, FDR corrected p <0.001. Dome, the domestic silkworm. Wild, the wild silkworm.

### *Knockout of* SP1 *by CRISPR/Cas9*

To explore the functional impact of *BmSp1* in silkworm domestication, we firstly investigated the biological role of this gene in the silkworm through CRISPR/Cas9 knocking out system. For single guide RNA (sgRNA) design, we selected highly specific targets in the first exon close to the translation starting site, namely, S1 and S2 (Fig 2A). We choose another site S3 close to the end of the first exon, more than 60 bp downstream of S1 and S2 (Fig 2A and Table 2), to obtain a potentially large fragment deletion by injecting the pool of three gRNAs. After mutation screening of the injected eggs (G0 generation), the gRNAs targeting the above three sites successfully guided DNA editing and generated a variety of mutation types, including 4–9 bp deletions or small insertions followed by a large deletion (Fig 2B).

**Table 2.**
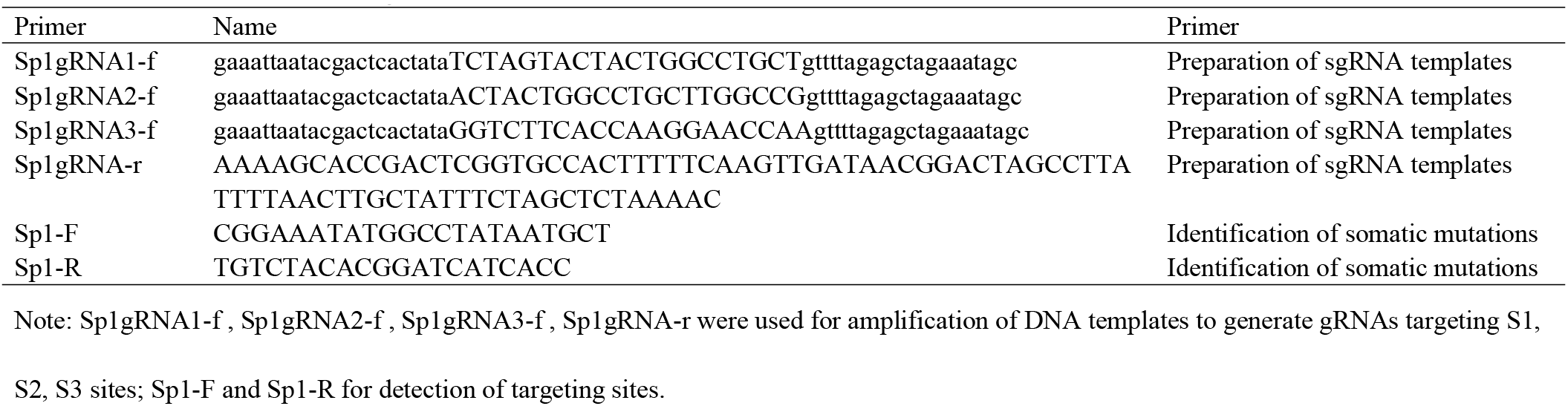
Primers used in this study

**Figure 2.**
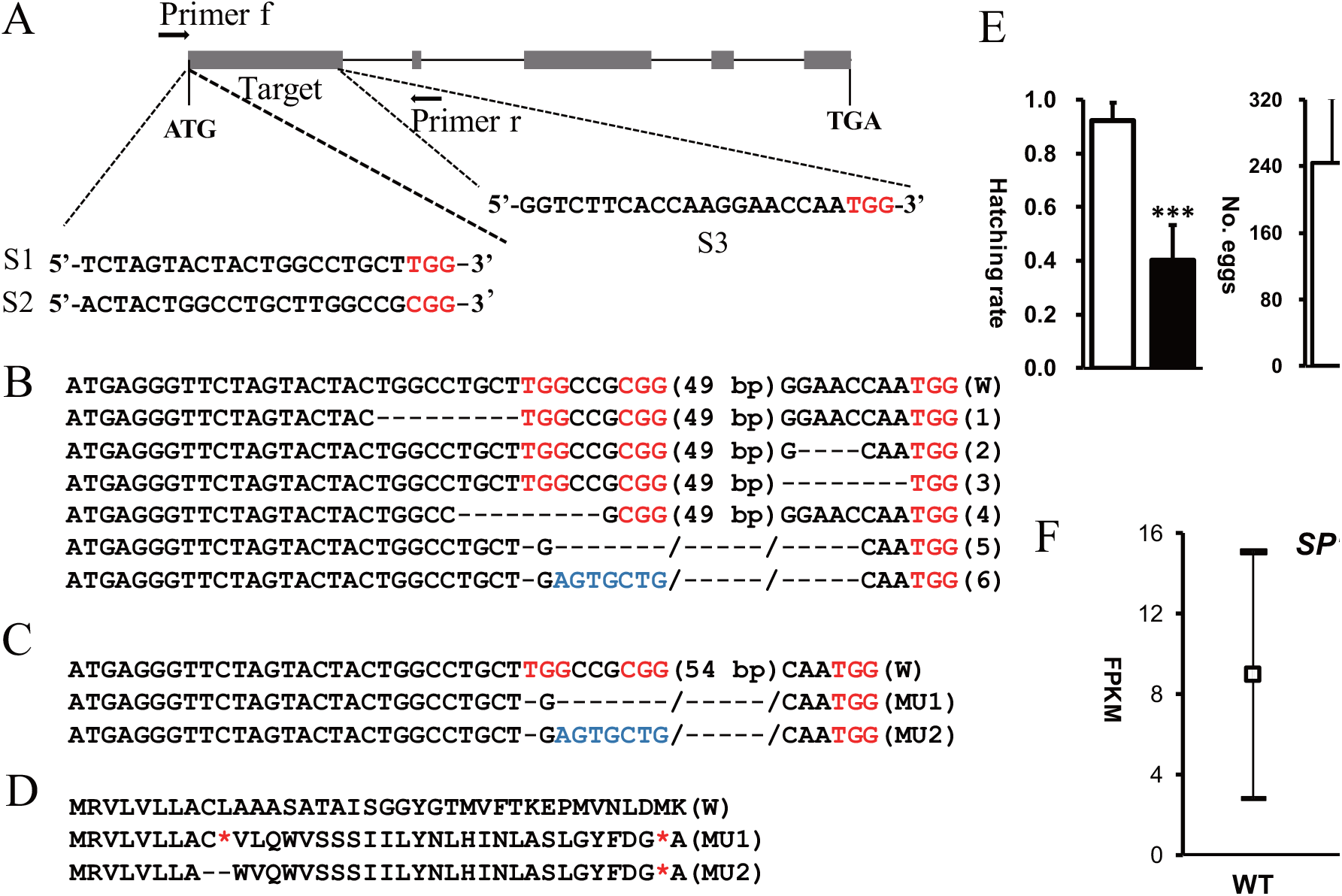
Cas9/sgRNA mediated gene editing of *SP1* in the silkworm. **(A)** Schematic diagram of sgRNA targeting sites. The five boxes indicate the five exons of *BmSp1*, and the black line with blocks represents the gene locus (blocks, exons; lines between blocks, introns). All sgRNA targeting sites are located on the first exon (S1, S2, and S3) and the PAM sequence is labeled in red. Sp1-F and Sp1-R were the primers used for mutation screening. **(B)** Various types of mutations in G0 injected embryos. Deletions are indicated by hyphens and insertions are shown in blue lowercase letters. The PAM sequence is in red. **(C)** Two types of mutations identified from homozygous mutant silkworms. W, wild type. MU1, *SP1* mutant type 1. There was an 8bp insertion followed by a 63 bp deletion in the *SP1* coding sequences; MU2, *SP1* mutant type 2. there was a 4 bp insertion followed by a 65 bp deletion. **(D)** Comparison of inferred amino acid sequences of homozygous mutant silkworms and the wild type. The missing amino acids are replaced with dashes. Premature stop codons are shown in red asterisk. **(E)** Phenotype assay of the mutants. Hatching rate of *SP1* mutants (MU1) decreased to about 51%. In total, 83 replicates of hatchings for MU1, 43 replicates for MU2 and 14 replicates for wild-type were used for statistical analysis. Number of egg produced, whole cocoon weight, pupa weight and cocoon layer weight, and pupa weight/whole cocoon weight ratio of mutants and wild-type silkworms, indicating no significant differences. In total, 32 replicates for mutants and 11 replicates for wild-type were used. Error bar: SD. **(F)** Expression level indicated as FPKM of *SP1* and *Vg* in the ova of domestic and wild silkworms. The three data pots indicate the highest value with 95% confidence, the average and the lowest value with 95% confidence, respectively. *, ** and *** represented significant differences at the 0.05, 0.01, 0.001 level (t-test).

By mutation screening of the exuviae of the fifth instar larva in the G0 cocoon, we successfully identified 26 mosaic mutant G0 moths. We then generated pairwise crosses of these G0 mutants with similar mutant genotypes for the G1 populations. After mutation screening of the G1 eggs, we selected two populations with large deletions for further feeding and mutation screening (see Material and methods).

Finally, in the G2 generation we obtained two types of homozygous mutants, i.e., MU1 and MU2 (Fig 2C). As to MU1, there was an 8 bp insertion followed by a 63 bp deletion in the *BmSP1* coding sequences. As to MU2, there was a 4 bp insertion followed by a 65 bp deletion. The mutations occurred at +29 bp of the first *SP1* exon in MU1 and MU2, respectively (Fig 2C), resulting in reading frame shift mutations and severe premature termination close to the translation starting site, with stop signals at +10 aa and +37 aa of theSP1 protein (Fig 2D).

### *Both* SP1 *mutants and the wild silkworm had decreased hatching rate and Vg expression*

We selected and maintained MU1 population for assay on phenotypes related to reproduction and metamorphosis, such as number of eggs, hatching rate, pupa weight, and cocoon weight. Compared with the wild-type, which showed hatching rates of about 90%, the hatching rates of *SP1* mutants were dramatically decreased, with a mean egg hatching rate of about 40% (Fig 2E), whereas the number of eggs produced was not obviously affected (Fig 2E), neither did the whole pupa weight or cocoon shell weight (Fig 2E). Given that the data were obtained from large replicates (126 replicates for hatchability assay and 320 replicates for pupa and cocoon weight), the results are convincible. Loss-of-function mutation resulted in significant decreased expression of *SP1* and *Vg* in the ova, based on the RNA-seq data (Fig 2F).

Given that knockout of *SP1* caused reduced hatching rate (Fig 2E) and that expression of *SP1* in the ova of wild silkworm were significantly lower than that of domestic one, we further suspected that artificial selection on *SP1* might improve silkworm hatching rate during domestication. Supporting this hypothesis, we found that hatching rates of the wild silkworm were generally lower compared with that of the domestic one and, similar to *SP1* mutant, no obvious differences was detected in egg production between the wild and the domestic silkworm (Fig 3A). The lower hatching rate of wild silkworm was also reported in other studies [21, 22]. Meantime, we also found that similar to SP1 mutants, expression of *Vg* in the ova of the wild silkworm was drastically lower compared with domestic one (Fig 3B). These results suggested that in the silkworm *SP1* may positively affect expression of *Vg* of ova and contribute to silkworm egg development. Promotion of *SP1* expression in the domestic silkworm thus results in the correspondingly up-regulation of *Vg*, which further contributes to increased hatchability during silkworm domestication.

**Figure 3.**
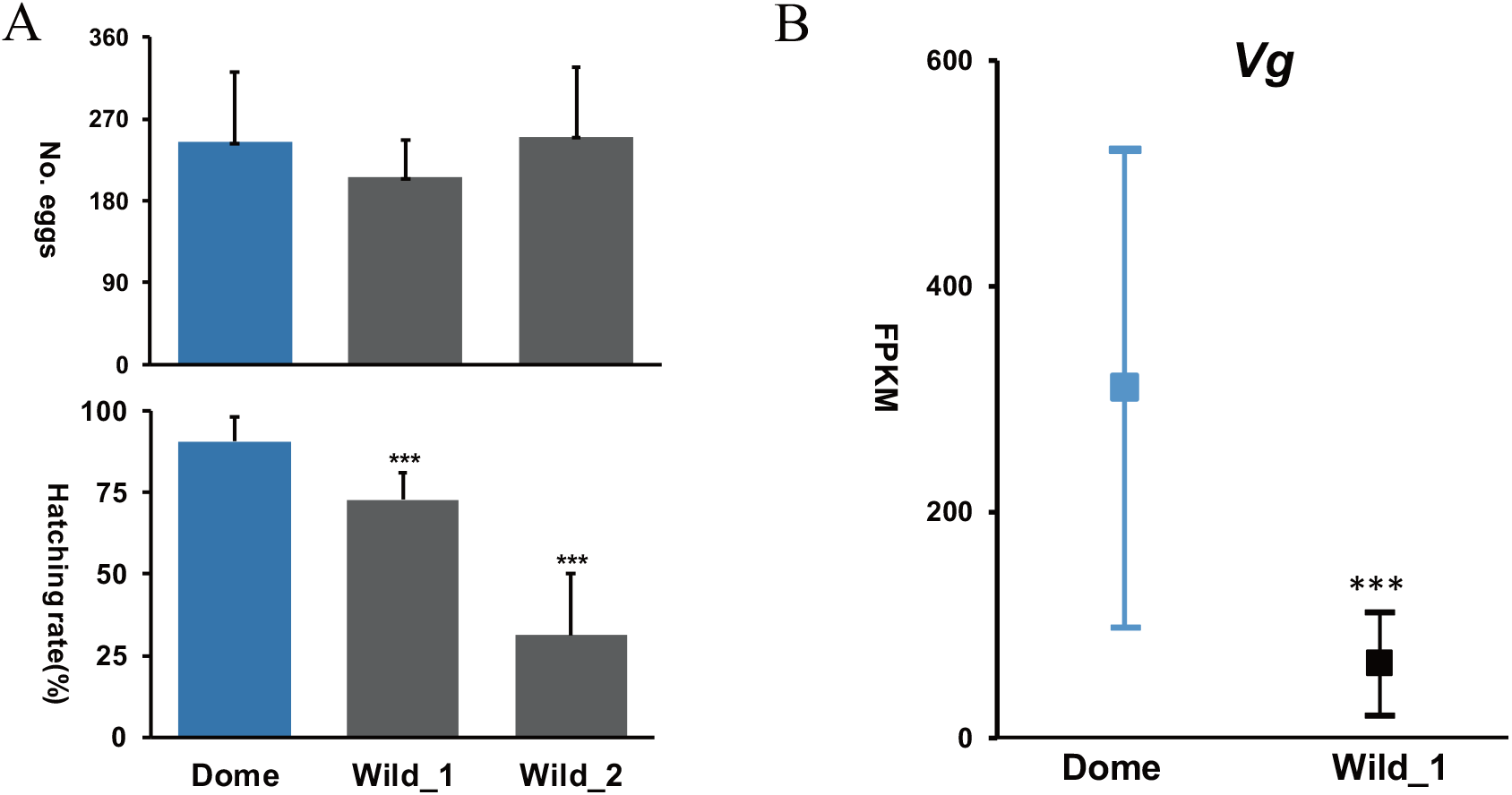
The artificial selection on *SP1* might improve silkworm hatching rate during domestication. **(A)** The egg production and hatching rates in the domestic group and wild silkworms. Error bar: SD. Wild-1 and Wild-2 indicated the results from two repeat times **(B)** Expression level indicated as FPKM of *Vg* in the ova of domestic and wild silkworms. The three data pots indicate the highest value with 95% confidence, the average and the lowest value with 95% confidence, respectively. *, ** and *** represented significant differences at the 0.05, 0.01, 0.001 level.

### Genes involved in egg development and eggshell formation were both repressed in SP1 mutant and the wild silkworm

In order to further exploring the regulation network and possible molecular mechanism of female specific *SP1* on egg hatchability, we generated comprehensively Ova comparative transcriptome analyses between the wild type and the mutant, as well as domestic and wild silkworm (*Bombyx mandarina*), with 4.87~9.15 Gb RNA-seq data for each sample (S1 Table). Totally, there were 561 genes identified as differential expressed genes (DEGs) in the SP1knockout mutants (MU1) compared to wild-type silkworm, with down-regulated genes (341) significantly more than up-regulated ones (220) (p=0.0003, Chi-squared test with Yates’ continuity correction) (Fig. 4A and S2 Table). As expected, there are much more differential expressed genes (2882) between the wild and domestic silkworm, since wild silkworm are much more genetically and phenotypically different from the domestic one, compared with the silkworm mutant. It is interesting that, down-regulated genes (1761) were also significantly more than up-regulated (1121) ones (p=2.2e-16, Chi-squared test with Yates’ continuity correction) (Fig 3A and S3 Table). The results suggested that transcriptome repression in ova might be an output of SP1 depletion, in both the SP1-KO mutant (Fig 2F and Fig 4A) and in the wild silkworm (Fig 1E and Fig 4A).

**Figure 4.**
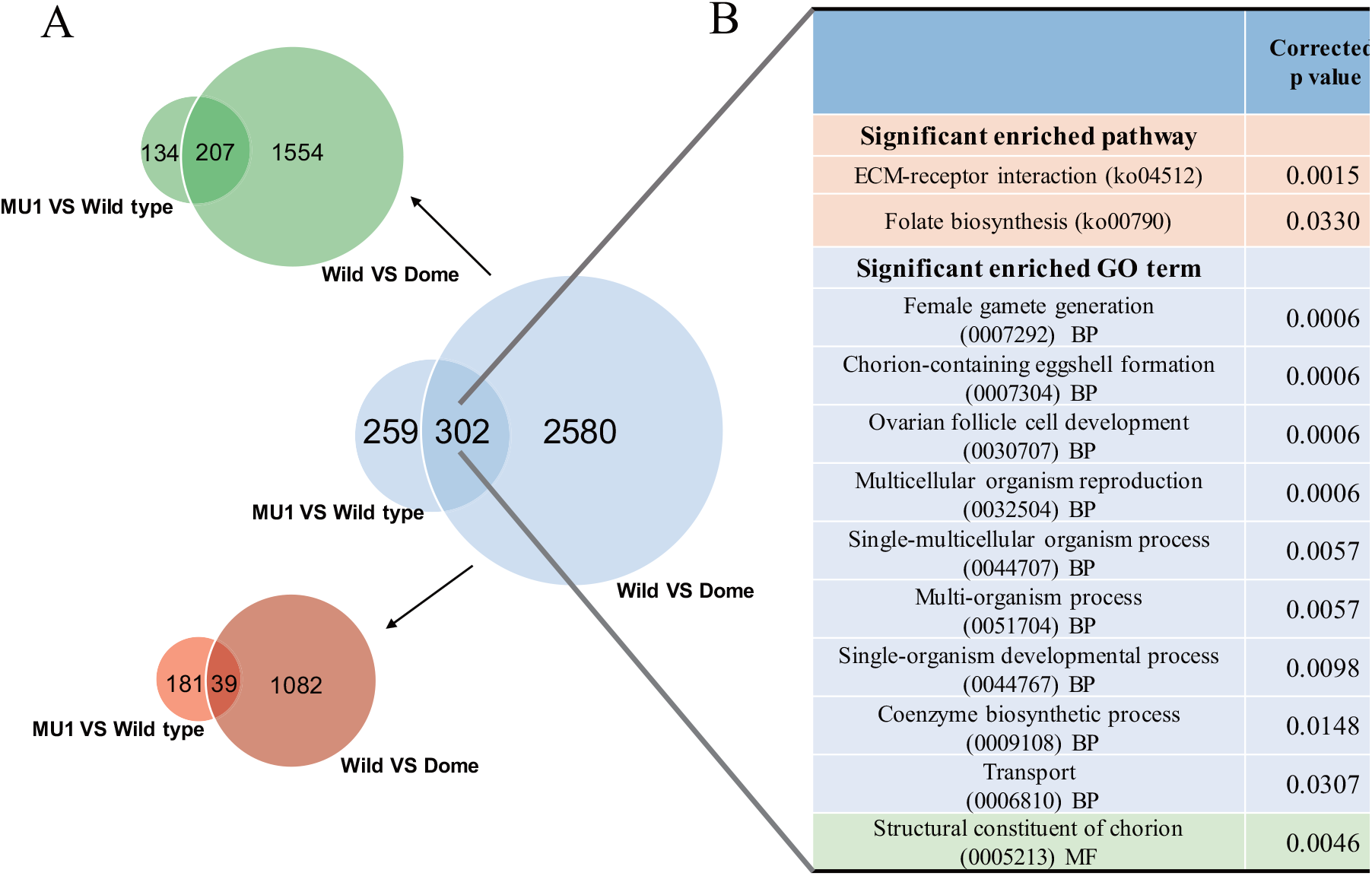
Differential expressed genes (DEGs) and functional enrichment analysis of the common genes from DEGs of two comparisons, i.e., from mutant VS the wild type, and from the wild and the dometic silworm. **(A)** Venn diagram of the DEGs between the wild type and the mutant, as well as domestic and wild silkworm (*Bombyx mandarina*). The blue venn diagram represents the total differentially expressed gene, The green venn diagram and the red venn diagram represent down-regulated differentially expressed and up-regulated differentially expressed, respectively. **(B)** Functional enrichment analysis of common genes shared in the two sets of DEGs. Corrected p-value: p-value in hypergeometric test after FDR correction. All non-redundant terms that had Corrected p-value < 0.05 is shown.

We identified 302 common genes shared in the two sets of DEGs. KEGG enrichment analysis indicated that these common differential expressed genes were significantly enriched in pathways related to cell proliferation, such as ECM-receptor interaction and folate biosynthesis. GO enrichment analysis indicated that they were enriched in reproduction related biological processes such as ovarian follicle cell development, chorion-containing eggshell formation (Fig 4B) and these genes was also enriched in the molecular function of structural constituent of chorion (Fig 4B). Factually they are all annotated as chorion proteins, including 4 chorion class CB protein M5H4-like genes (*BGIBMGA003248, BGIBMGA009720, BGIBMGA009719, BGIBMGA009715*), a chorion class B protein PC10 (BGIBMGA009721) and a Chorion 1 domain containing gene (*BGIBMGA005877*). All these chorion like genes are down regulated except *BGIBMGA009720* (Fig 4B, Table S2, S3). Genes in ECM-receptor interaction pathway includes collagens and integrins (S2 Fig) and those in folate biosynthesis includes folylpolyglutamate synthase, involved in 7,8-Dihydrofolate (DHF) and 5,6,7,8-Tetrahydrofolate (THF) which are substrates for subsequent one carbon pool mediated by folate (S3 Fig). Extend to all the enriched genes, it is notable that they were mostly down-regulated, in both the SP1 mutant and the wild silkworm (Fig 4B). This results suggested that the common influence of repression of *SP1* in both the SP1 mutant and the wild silkworm at transcriptome level is on genes or pathways involved in reproduction, such as ovarian follicle cell development or proliferation, and eggshell formation.

We further generated enrichment analyses on DEGs in the two sets of comparison independently and observed consistent pattern (Table 3 and 4). Functional enrichment analysis of DGEs between wild-type silkworm and *SP1* mutant silkworm reveals significantly enriched the KEGG pathway “ECM-receptor interaction” and the GO terms, including ovarian follicle cell development and eggshell formation process, functioning as structural constituent of chorion (Table 3) and the related genes are mostly down-regulated. Consistently, These GO items were also in the top rank with the lowest p value when analyzing the DEGs between wild and domestic silkworm (Table 4). The involved genes in these KEGG or GO terms were nearly all down-regulated in the wild silkworm (Table 4). These results further supported that in the wild silkworm, repression of *SP1* may result in suppressed expression of ovarian follicle cell development and eggshell formation process.

**Table 3.**
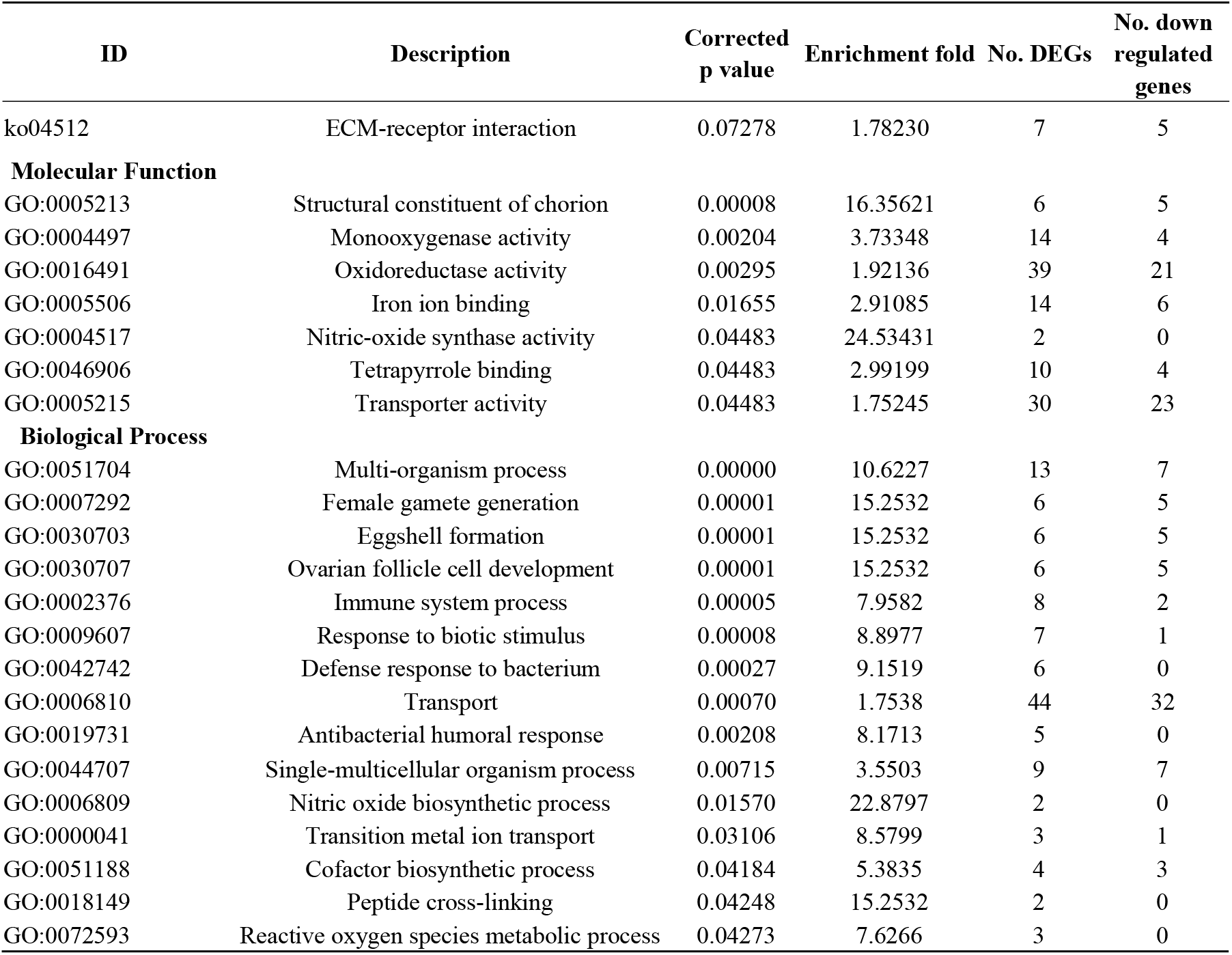
Functional Enrichment of differential expressed genes between the loss-of-funtion *SP1* mutants and the wild-type domestic silkworms.

| **ID** | **Description** | **Corrected**<br>**p value** | **Enrichment fold** | **No. DEGs** | **No. down**<br>**regulated**<br>**genes** |
| --- | --- | --- | --- | --- | --- |
| ko04512 | ECM-receptor interaction | 0.07278 | 1.78230 | 7 | 5 |
| **Molecular Function** |  |  |  |  |  |
| GO:0005213 | Structural constituent of chorion | 0.00008 | 16.35621 | 6 | 5 |
| GO:0004497 | Monooxygenase activity | 0.00204 | 3.73348 | 14 | 4 |
| GO:0016491 | Oxidoreductase activity | 0.00295 | 1.92136 | 39 | 21 |
| GO:0005506 | Iron ion binding | 0.01655 | 2.91085 | 14 | 6 |
| GO:0004517 | Nitric-oxide synthase activity | 0.04483 | 24.53431 | 2 | 0 |
| GO:0046906 | Tetrapyrrole binding | 0.04483 | 2.99199 | 10 | 4 |
| GO:0005215 | Transporter activity | 0.04483 | 1.75245 | 30 | 23 |
| **Biological Process** |  |  |  |  |  |
| GO:0051704 | Multi-organism process | 0.00000 | 10.6227 | 13 | 7 |
| GO:0007292 | Female gamete generation | 0.00001 | 15.2532 | 6 | 5 |
| GO:0030703 | Eggshell formation | 0.00001 | 15.2532 | 6 | 5 |
| GO:0030707 | Ovarian follicle cell development | 0.00001 | 15.2532 | 6 | 5 |
| GO:0002376 | Immune system process | 0.00005 | 7.9582 | 8 | 2 |
| GO:0009607 | Response to biotic stimulus | 0.00008 | 8.8977 | 7 | 1 |
| GO:0042742 | Defense response to bacterium | 0.00027 | 9.1519 | 6 | 0 |
| GO:0006810 | Transport | 0.00070 | 1.7538 | 44 | 32 |
| GO:0019731 | Antibacterial humoral response | 0.00208 | 8.1713 | 5 | 0 |
| GO:0044707 | Single-multicellular organism process | 0.00715 | 3.5503 | 9 | 7 |
| GO:0006809 | Nitric oxide biosynthetic process | 0.01570 | 22.8797 | 2 | 0 |
| GO:0000041 | Transition metal ion transport | 0.03106 | 8.5799 | 3 | 1 |
| GO:0051188 | Cofactor biosynthetic process | 0.04184 | 5.3835 | 4 | 3 |
| GO:0018149 | Peptide cross-linking | 0.04248 | 15.2532 | 2 | 0 |
| GO:0072593 | Reactive oxygen species metabolic process | 0.04273 | 7.6266 | 3 | 0 |

**Table 4.** Functional enrichment of differential expressed genes between the wild and domestic silkworms.

| ID | Description | Corrected<br>p value | Enrichment<br>fold | No.<br>DEGs | No. down<br>regulated genes |
| --- | --- | --- | --- | --- | --- |
| ko03010 | Ribosome | 0.00001 | 2.08066 | 50 | 49 |
| **Molecular**<br>**Function** |  |  |  |  |  |
| GO:0003735 | Structural constituent of ribosome | 0.00003 | 2.12234 | 44 | 42 |
| **Biological Process** |  |  |  |  |  |
| GO:0007292 | Female gamete generation | 0.12304 | 2.99379 | 6 | 5 |
| GO:0007304 | Chorion-containing eggshell formation | 0.12304 | 2.99379 | 6 | 5 |
| GO:0046700 | Heterocycle catabolic process | 0.12304 | 2.56610 | 8 | 5 |
| GO:0009653 | Anatomical structure morphogenesis | 0.12304 | 2.99379 | 6 | 5 |
| GO:0019953 | Sexual reproduction | 0.12304 | 2.99379 | 6 | 5 |
| GO:0030707 | Ovarian follicle cell development | 0.12304 | 2.99379 | 6 | 5 |
| GO:1901361 | Organic cyclic compound catabolic process | 0.12304 | 2.69441 | 9 | 6 |
| GO:1901564 | Organonitrogen compound metabolic process | 0.12304 | 1.29411 | 117 | 98 |
| GO:0043603 | Cellular amide metabolic process | 0.12304 | 1.50499 | 62 | 57 |
| GO:0043604 | Amide biosynthetic process | 0.12304 | 1.48862 | 60 | 55 |
| GO:0043043 | Peptide biosynthetic process | 0.12304 | 1.42885 | 56 | 52 |

### Ribosome proteins and genes in amide and peptide biosynthetic processes were also repressed in the wild silkworm

DEGs between wild and domestic silkworms were significantly in enriched in function of structural constituent of ribosome. The related genes are mostly ribosome proteins (S4 Fig). We also noted that the related biological progresses, such as amide and peptide biosynthetic processes was also in the top rank with the lowest p value (Table 4). The related genes were also mostly down-regulated in the wild silkworm. However, during domestication, there might be other factors that contributes to improved hatchability, that is, improved amide and peptide biosynthesis and activated ribosome activities in the ovarian.

## Discussion

Nitrogen resource is very important for silkworm domestication. The domestic silkworm tend to have efficient utilization of nitrogen resources for yielding protein outputs to adapt to human-preference. Here in this study, we discovered that artificial selection could directly act on nitrogen resource gene, i.e, storage protein1 (*SP1*), for improved silkworm hatchability. SPs are also of target loci of breeding in crops [23]. However, in the crops, human could directly benefit from the nutrient of these improved SPs [15] whereas in the silkworm, the benefits of SPs are increased silkworm reproductive capacity.

Among all the SPs identified, SP1 is quite diverged and somewhat unique from the others, in terms of both genomic location and phylogenetic position. Similar pattern was also observed in other Lepidoptera species, such as tobacco hornworm, *Manduca sexta* [24], suggesting that *SP1* might evolve dependently while the other type of *SPs* might have experienced duplication during Lepidoptera evolution. Methionine-rich SP1 seems to be of special interest, since methionine is reported to be an important amino acid for the tradeoff between growth and reproduction [25]. In *Drosophila*, dietary methionine restriction extends lifespan [25], while in grasshoppers, a reduced reproduction-induced increase in expression methionine-rich protein occurred during life extension [26]. Similarly, in the beet armyworm, silencing of Sp1 by RNA interference (RNAi) decreases larval survival, which indicate the role of the methionine-rich SP in growth and metamorphosis[12]. We therefore added an new evidence that different to grasshopper [26] and the beet armyworm[12], but similar to *Drosophila*[25], silkworm methionine-rich SP1 functions in reproduction process but not obviously affect growth.

Given that in those cocoon-producing silk moths, another nitrogen utilization system such as gluminate/glutamine cycle system were reported to be vital in metamorphosis silk-cocoon production [1, 4, 27], we suspect that strategy of nitrogen resource allocation via storage proteins may be diverged or modified during Lepidoptera insect evolution. Here in the silkworm, function of SP1 limited to influencing egg hatching rate. Artificial selection acted only on *SP1*, again suggesting functional importance of SP1 rather than the other SPs, for human-preferred domestication traits, i.e, increased hatchability.

Ova comparative transcrtiome analyses further illustrated a frame of regulation network of SP1 on hatchability. Vitellogenin(Vg), chorion proteins, structural component proteins in the extracellular matrix (ECM)-interaction pathway such as collagen and integrins, and synthetase in folate biosynthesis are all generally repressed in both the *SP1* mutant and the wild silkworm. Thus, artificial selection acting on *SP1* for increased hatchability, possibly thorough direct or indirect influence on those genes, pathway or biological processes. Vg is the main nutrition for silkworm egg formation and embryonic development. It appears and accumulates at the stage when SP1 rapidly declined and disappeared in the fat body shortly before adult emergence [16, 28]. SP1 may supplies amino acids for synthesis of Vg, as previously reported in *Plutella xylostella* (Yaginuma & Ushizima, 1991). Chorion proteins are the major component of the silkworm eggshell that have the essential function of protecting the embryo from external agents during development while allowing gas exchange for respiration. Eggshell (chorion) is constructed by the ovarian follicle cells. The follicle cell epithelium surrounds the developing oocyte and in the absence of cell division synthesizes a multilayer ECM [29]. Eggshell ECM were usually linked by integrins, a family of transmembrane receptor proteins to the cytoskeleton of oocyte. Via a series of signal transduction, ECM-integrins functions in oocyte movement, differentiation, and proliferation [29]. Integrins was reported to function in formation of actin arrays in the egg cortex [30] and it is also involved in tracheole morphogenesis which is for respiration [31]. Folate is one of B-vitamin cofactors with and functions in transferring various single-carbon units. The key folylpolyglutamate synthase is involved in production of 7,8-Dihydrofolate (DHF) and 5,6,7,8-Tetrahydrofolate (THF) which are substrates for subsequent one carbon pool mediated by folate. In human, folate is used as a supplement by women during pregnancy to prevent neural tube defects (NTD) in the baby, and low levels in early pregnancy are believed to be the cause of more than half of babies born with neural tube defects, indicating its important role for fetal development [32]. In insects, folate also plays important roles in egg development. It could promote biosynthesis of nucleic acids in the ovaries, and evoke mitoses in cells of the collicular epithelium [33–35].

Notably, increased hatchability during domestication may not be solely attributed to increased expression of *SP1* and the associated downstream genes, given that artificial selection acts on hundreds of gene loci in silkworm genome [1, 36] and that ova comparative transcriptome between wild and domestic silkworm identified much more genes than that between SP1 mutant and wild type silkworm. We factually observed significantly enriched pathway and structural constituent of ribosome, the protein translation machinery and involved biological processes in amide and peptide biosynthesis, are generally lower expressed in the wild silkworm compared with domestic silkworm (Table 4). These results again supported the importance of nitrogen and amino acid in silkworm domestication, not only for silkworm protein output [1], but also for productivity.

Similar to other domesticates, hatchability of silkworm eggs directly determine quantity of offspring, and thus it is an important productivity trait human favorably selected during domestication. Based on the results and the discussion above, we proposed that artificial selection, on one hand favored higher expression of *SP1* in the domestic silkworm, which would subsequently up-regulate the genes or pathways vital for egg development and eggshell formation. On the other hand, artificial selection consistently favored activated ribosome activities and improved amide and peptide biosynthesis and in the ova, as it might act in the silk gland for increased silk-cocoon yield [1]. As an output, domestic silkworm demonstrates improved increase egg hatchability compared with it wild ancestor.

## Materials and Methods

### Silkworm strains

A multivoltine silkworm strain, Nistari, was used in all experiments. Larvae were reared on fresh mulberry leaves under standard conditions at 25°C. The wild silkworms were collected in Zhejiang province, China and maintained as laboratory population in our lab.

### Silkworm genomic data resource

The silkworm *Bombyx mori* reference genome and related data used for searching and screening of artificial selection signature on silkworm SPs is our updated version (https://doi.org/10.5061/dryad.fn82qp6); those for RNA-seq data analyses were obtained from Ensemble database (http://metazoa.ensembl.org/Bombyx_mori/Info/Index).

### Genomic search of B.mori SPs and Phylogenetic analysis of SPs

The reference sequences of *B. mori* storage proteins (SP1, SP2, SSP2 and SP3) were retrieved from the NCBI GenBank. These sequences were used as quires searching for homologs in the *B. mori* genome by tblastn with e-value <10^-7^ Other insect homologs of the silkworm SPs were searched in GenBank (https://blast.ncbi.nlm.nih.gov/) by BLASTP with an e-value <10^-7^ We selected sequences from several representative Lepidoptera species and *Drosophila melanogaster* as candidate proteins for further analyses. The sequences of the SP1 homologs were aligned using MEGA 6.0 software [37]. A gene tree was constructed using The Bayes tree was constructed by MrBayes-3.1.2 with GTR + gamma substitution model. The gene-ration number was set as 1000000 and the first 25% was set as burn-in. Other parameters were set as default.

### Molecular selection of Sp1 in domesticated and wild silkworm populations

Based on the available whole genomic Single nuclear polymorphic data (SNP) of domesticated and wild silkworm populations [1], (https://doi.org/10.5061/dryad.fn82qp6), we screened the selection signatures of the silkworm *SPs*, according to Xiang et al’s pipeline [1]. Allelic frequency and SNP annotation was calculated by in-house Perl scripts.

### Design of sgRNA target and in vitro synthesis of sgRNA and Cas9 mRNA

The 20 bp sgRNA targets immediately upstream of PAM were designed by the online platform CRISPRdirect (http://criSpr.dbcls.jp/)[38]. The sgRNA DNA template was synthesized by PCR, with Q5^®^ High-Fidelity DNA Polymerase (NEB, USA). The PCR conditions were 98°C for 2 min, 35 cycles of 94°C for 10 s, 60°C for 30 s, and 72°C for 30 min, followed by a final extension period of 72°C for 7 min. The sgRNA were synthesized based on the DNA template *in vitro* using a MAXIscript^®^ T7 kit (Ambion, Austin, TX, USA) according to the manufacturer’s instructions. The Cas9 construct was a kind gift provided by the Shanghai Institute of Plant Physiology and Ecology (Shanghai, China). The Cas9 vector was pre-linearized with the NotI-HF^®^ restriction enzyme (NEB, USA). The Cas9 mRNA was synthesized *in vitro* with a mMESSAGE mMACHINE^®^ T7 kit (Ambion, Austin, TX, USA) according to the manufacturer’s instructions. All related primers are shown in Table 2.

### Microinjection of Cas9/gRNAs

Fertilized eggs were collected within 1 h after oviposition and microinjection was within 4 h. The Cas9-coding mRNA (500 ng/μL) and total gRNAs (500 ng/μL) were mixed and injected into the preblastoderm Nistari embryos (about 8 nl/egg) using a micro-injector (FemtoJet^®^, Germany), according to standard protocols (Tamura, 2007). The injected eggs were then incubated at 25°C for 9–10 d until hatching.

### Cas9/gRNAs-mediated mutation screening and analyses of germline mutation frequency

To calculate the efficiency of Cas9/gRNAs-mediated gene mutation in the injected generation (G0), we collected ~10% of the eggs (64 out of 600) 5 d after injection to extract genomic DNA for PCR, with primers Sp1-F and Sp1-R (Table 2). The amplified fragments were cloned into a pMD^TM^19-T simple vector (Takara, Japan) and sequenced to determine mutation type and mutagenic efficiency.

When the injected G0 silkworms pupated, we collected silkworm exuviae from fifth instar larvae in each cocoon. Individual DNA was then extracted Genomic DNA was extracted using a TIANamp Blood DNA Kit (Tian gen Biotech, Beijing) according to the manufacturer’s instructions. Individual mutation screening was generated with PCR with 94°C for 2 min, 35 cycles of 94°C for 30 s, 57°C for 30 s, and 72°C for 45 s, followed by a final extension period of 72°C for 5 min. PCR products were cloned to pMD^TM^19-T simple vector (Takara, Japan) and sequenced.

### Homozygous mutant screening and mutation effect on inferred protein

Mosaic mutant moths were obtained from the above mutation screening of exuviae DNA from fifth instar larvae. Moths with the same mutation site were pairwise crossed with each other to acquire G1 offspring. About 7 d after the G1 eggs were laid, we collected ~30 eggs of each offspring population from one parental pair and pooled them to extract genomic DNA for mutation screening by PCR. The amplified fragments were cloned into a pMD^TM^19-T simple vector (Takara, Japan) and sequenced to determine the exact mutation type. Two G1 offspring populations with large deletions in *BmSp1* were selected for further breeding. At the pupa stage, 20 randomly selected individuals within each population were subjected to mutation screening of exuviae DNA. Homozygous mutant moths with the same identified mutant genotype were crossed for G2 offspring. Mutation effects on proteins were evaluated using MEGA 6/0 software[37] by codon alignment of the wild type and the mutant.

### Phenotypic assay

On the fourth day of pupation (P4), the silkworms had successfully pupated. We weighed every 10 individuals as a group, recorded the whole cocoon weight, pupa weight, and cocoon shell weight, respectively, and calculated the ratio of pupa weight to whole cocoon weight. In total, 32 replicates for the mutants and 11 replicates for the wild-type silkworms were set, respectively. Offspring of the homozygous mutants and wild-type silkworms were incubated at 25°C for 9–10 d until hatching. Number of Egg produced and egg hatching rate were determined. Eighty-three, forty-three, and fourteen replicates were set for the two mutant (SP1-MU1, SP1-MU2) and wild-type populations.

Number of eggs produced and the hatching rate of the wild silkworm were also recorded. Ten replicated was set repeated two times.

### Ova dissection, total RNA isolation and RNA-seq

Ova from virgin moth of the domestic wild type silkworm, SP1 mutant and the wild type were dissected and used to extract total RNA with three replicates. Total RNA were isolated using TRIzol (Invitrogen). For each sample, RNA were sent to Novogene Bioinformatics Institute (Beijing, China) for cDNA library construction and RNA-seq. Six cDNA libraries were sequenced by Illumina Hiseq 2500 (Illumina, San Diego, CA, USA) with 125 bp paired-end reads according to the manufacturer’s instructions.

### Analyses on RNA-seq data

Raw data were filtered with the following criteria: (1) reads with ≥ 10% unidentified nucleotides (N); (2) reads with > 10 nt aligned to the adapter, allowing ≤ 10% mismatches; and (3) reads with > 50% bases having phred quality < 5. The clean data were mapped to the *Bombyx mori* reference genome using Tophat with 2 nt fault tolerance and analyzed using Cufflinks [39]. The expression value of each gene was calculated and normalized using fragments per kilobase of exon per million reads mapped (FPKM) [39]. In order to identify differentially expressed genes, Cuffdiff was used to perform pairwise comparisons between wild-typed and SP1 mutant samples, as well as the wild and domestic silkworm, respectively, with corrected *P*-value of 0.05.<5 and Log2-fold change>1.

KEGG and GO enrichment analyses of differentially expressed genes were performed with an online platform (http://www.omicshare.com/tools/), using all the annotated genes in *Bombyx mori* as background.

### Data availability

RNA-seq data were deposited in the NCBI Short Read Archive database under the accession SAMN09700389-SAMN09700397.

## Acknowledgements

We thank Anli Chen, Muwang Li and Lang You for their kind helps in silkworm maintaining, and Anjiang Tan for his kindly providing the Cas9 vector. We thank Shuai Zhan for his help in genomic data analyses and mining. This work was supported by a National Basic Research Program of China (No. 2013CB835204) and a grant of National Natural Science Foundation of China (No. 31371286).

## Supporting Information Caption

**S1 Fig. SNPs between wild and domestic silkworm groups that cause non-synonymous mutations of SP1. (A).** Amino acid alignment of SP1 protein of the domestic silkworm (*B.mori*) and the deduced SP1 protein sequence of the wild silkworm (*B. mandarina*) by inferred SNP data. **(B)**. Allelic frequencies of the 11 non-synonymous mutations in the wild (the left column) and domestic silkworm (the right column). Blue, reference allele; Black, alternative allele. The 11 mutations of amino acid were labelled with numerals in red.

**S2 Fig. Scheme of ECM-receptor interaction pathway.** The shared DEGs of ova in two sets of comparison (SP1 mutant vs wild type and wild vs domestic silkworm) are indicated by red frames.

**S3 Fig. Scheme of folate biosynthesis pathway.** The shared DEGs of ova in two sets of comparison (SP1 mutant vs wild type and wild vs domestic silkworm) are indicated by red frames.

**S4 Fig. Scheme of Ribosome pathway.** The DEGs of ova between the wild and domestic silkworm are indicated by red frames.

**S1 Table. Summary of RNA-seq data.**

**S2 Table. Information of the differentially expressed genes between SP1 mutant (MU1) and the wild type (WT) domestic silkworm.**

**S3 Table. Information of the differentially expressed genes between the wild and the domestic (Dome) silkworms.**

